# A genetic variant protective against severe COVID-19 is inherited from Neandertals

**DOI:** 10.1101/2020.10.05.327197

**Authors:** Hugo Zeberg, Svante Pääbo

**Affiliations:** Max Planck Institute for Evolutionary Anthropology, Deutscher Platz 6, D-04103 Leipzig, Germany; Department of Neuroscience, Karolinska Institutet, SE-17177 Stockholm, Sweden; Okinawa Institute of Science and Technology, Onna-son, Okinawa 904-0495, Japan

## Abstract

It was recently shown that the major genetic risk factor associated with becoming severely ill with COVID-19 when infected by SARS-CoV-2 is inherited from Neandertals. Thanks to new genetic association studies additional risk factors are now being discovered. Using data from a recent genome-wide associations from the Genetics of Mortality in Critical Care (GenOMICC) consortium, we show that a haplotype at a region associated with requiring intensive care is inherited from Neandertals. It encodes proteins that activate enzymes that are important during infections with RNA viruses. As compared to the previously described Neandertal risk haplotype, this Neandertal haplotype is protective against severe COVID-19, is of more moderate effect, and is found at substantial frequencies in all regions of the world outside Africa.

## Main text

Neandertals and Denisovans are archaic hominin groups that became extinct about 40,000 years ago (Higham *et al*. 2014). They have a biological impact on human physiology today through genetic contributions to modern human populations that occurred during the last tens of thousands of years of their existence (*e.g*., Simonti *et al*. 2016, Dannemann and Kelso 2017). Several of these contributions affect genes involved in the immune system (*e.g*., Laurent *et al*. 2011, Quach *et al*. 2016). In particular, variants at several loci containing genes involved in innate immunity come from Neandertals and Denisovans, for example toll-like receptor gene variants which decrease the susceptibility to *Helicobacter pylori* infections and increase the risk for allergies (Dannemann *et al*. 2016).

Recently, it was shown that a region on chromosome 3 harbors the major genetic risk locus for becoming critically ill upon infection with the novel coronavirus SARS-CoV-2 (Ellinghaus *et al.* 2020) and that the haplotype that confers the risk was contributed by Neandertals to modern humans (Zeberg and Pääbo 2020). This haplotype reaches carrier frequencies of up to ~65% in South Asia whereas it is almost absent in East Asia and occurs at ~16% in Europe. Thus, although this haplotype is detrimental for its carriers during the current pandemic, it has likely been beneficial in earlier times in South Asia, perhaps by conferring protection against other pathogens.

A new study from the GenOMICC consortium, which includes 2,244 critically ill COVID-19 patients and controls (Pairo-Castineira *et al*. 2020), identifies three new loci with genome-wide significant effects in addition to the risk locus on chromosome 3 located on chromosomes 12, 19 and 21. To test if any of these loci carry variants derived from Neandertals, we investigated if alleles of the index variants at these loci co-segregate in the population with alleles of single nucleotide polymorphisms (SNPs) that match three high-quality Neandertals genomes, while being absent in the genomes of 108 African Yoruba individuals (r^2^ > 0.75 in all genomes of the 1000 Genomes Project). No SNPs fulfilling these criteria were found in two of the novel loci whereas 43 such SNPs exists at the chromosome 12 locus.

To further investigate this locus, we used the round 4 (alpha) release of the COVID-19 Host Genetics Initiative (HGI) (covid19hg.org) to analyze 35 SNPs that are significantly associated with severe COVID-19 and have been scored in the archaic genomes (Table S1). A 60,000 to 80,000-year-old Neandertal from Chayrskaya Cave in southern Siberia (Mafessoni *et al*. 2020) carries the protective alleles at 34 of these SNPs in a homozygous form, while an approximately 50,000-year-old Neandertal from Vindija in Croatia (Prüfer *et al*. 2017) carries the protective alleles in a homozygous form at 33 SNPs. An approximately 120,000-year-old Neandertal from Denisova Cave in southern Siberia (Prüfer *et al*. 2014) carries protective alleles in homozygous form at 30 of the SNPs and in a heterozygous form at three of the SNPs. In contrast, only two of the protective alleles match the Denisovan genome (Meyer *et al*. 2012). Thus, for almost all SNPs with genome-wide significance on chromosome 12, the Neandertals carry the allele that decrease the risk of requiring hospitalization upon SARS-CoV-2 infection. In the GenOMICC study, the risk of carriers of the index variant of the haplotype (rs10735079, p=1.7e-8) of needing intensive care is reduced by ~22% (under the rare disease assumption, OR = 0.78, 95% confidence interval (CI): 0.71-0.85).

By identifying sets of Neandertals alleles that co-segregate in humans living today, we find a Neandertal-like segment of ~90 kilobases (kb) (chr12:113,345,168-113,435,449; *hg19*) which overlaps the genomic region harboring SNPs associated with severe COVID-19 (Fig 1; Methods). Genomic segments with similarity to Neandertal genomes may either derive from common ancestors of the two groups living about half a million years ago or be contributed by Neandertals to modern humans when the two groups met less than 100,000 years ago (Sankararaman *et al*. 2012). To distinguish these two alternatives, we use the recombination rate of the genomic region (0.80cM/Mb) (Kong *et al*. 2002), a generation time of 29 years (Langergraber *et al*. 2012), a split between Neandertals and modern humans of 550,000 years (Prüfer *et al*. 2014), interbreeding between the two groups ~50,000 years ago, and a published equation (Huerta-Sánchez *et al*. 2014) to test if a segment of this length may have survived since the common ancestor of the groups without being broken down by recombination that affect chromosomes in each generation. Under these assumptions, we exclude that it derives from the common ancestor (p = 1.1e-10) and conclude that this region entered the human gene pool from Neandertals. In agreement with this, a previous study (Mendez *et al*. 2013) has described gene flow from Neandertals in this region. We find that the index allele, *i.e*. the Neandertal allele with the greatest association with severe COVID-19 (rs4766664), occurs in all populations in Eurasia and Americas at frequencies that often reach and exceed 30% (Fig. 2).

**Figure 1.**
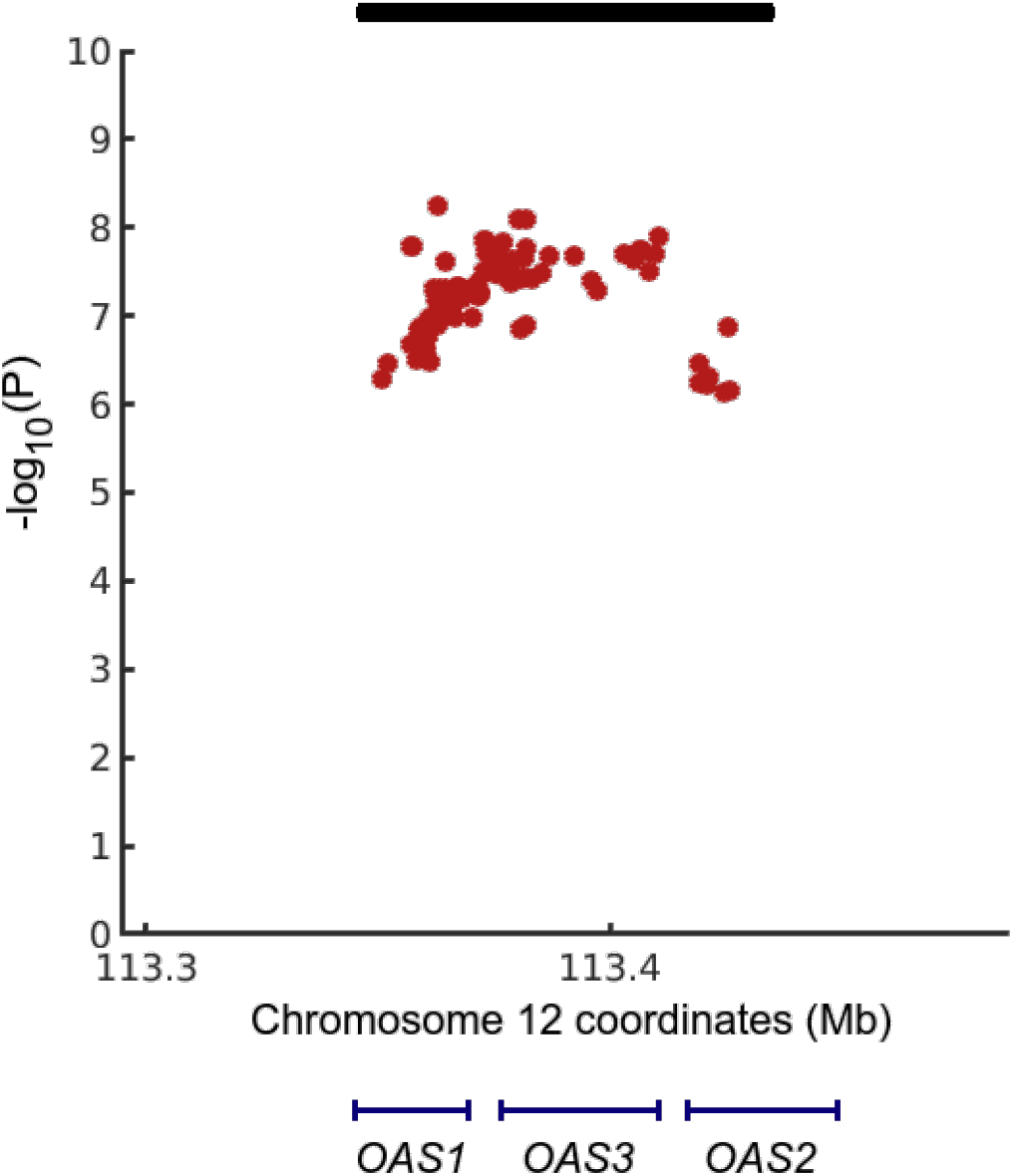
Genetic variants associated with COVID-19 hospitalization (p<1e-6) at the *OAS* locus. Black bar indicates a Neanderal haplotype defined by 66 diagnostic Neandertal variants in linkage disequilibrium in Europeans (r^2^>0.75, Methods, Fig. S1). Blue lines indicate the three *OAS* genes, all of which are transcribed from left to right. The *X*-axis gives *hg19* coordinates.

**Figure 2.**
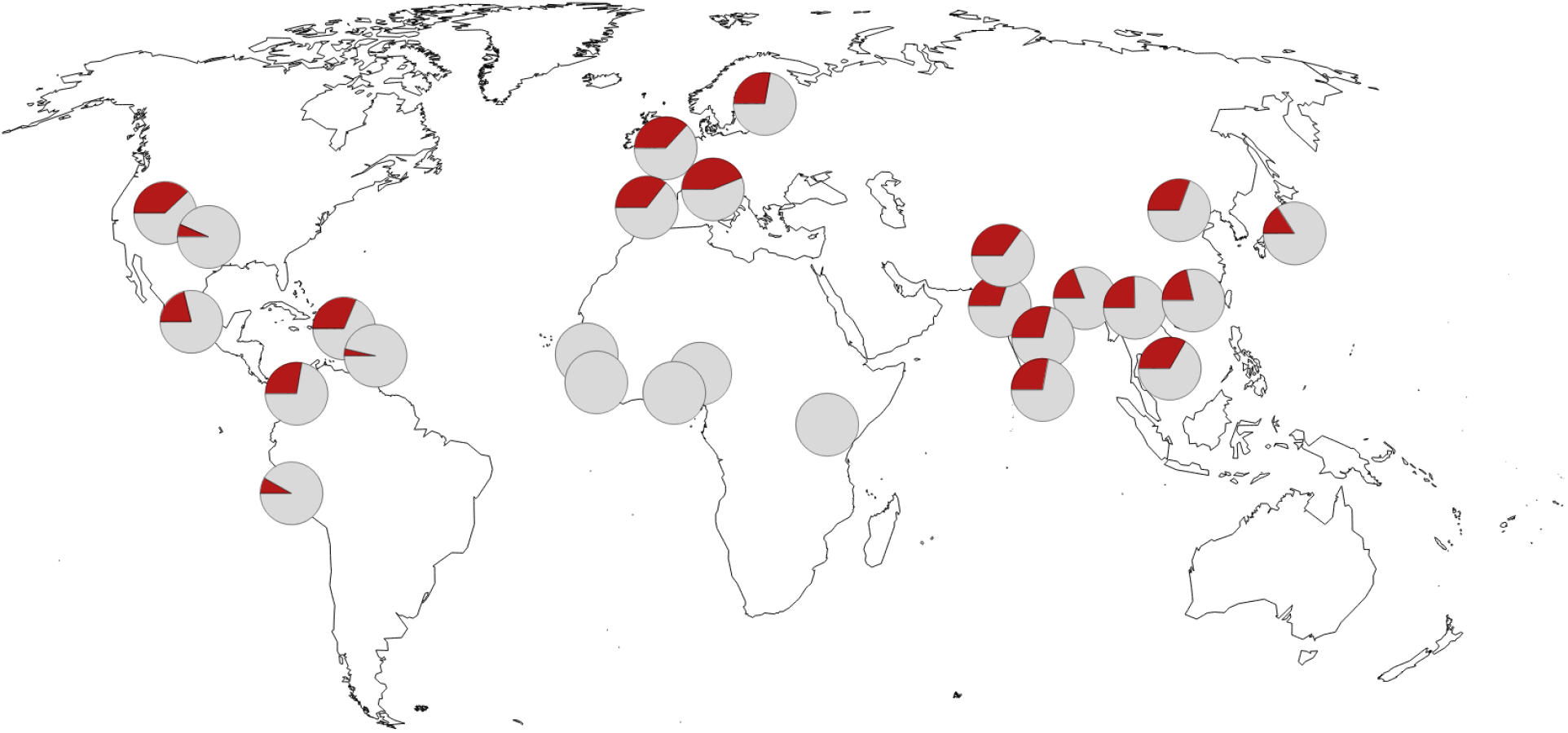
Geographic distribution of the allele indicative of the Neandertal haplotype protective against severe COVID-19. Pie charts indicate minor allele frequency at rs4766664. Frequency data from the 1000 Genomes Project. Map source data from OpenStreetMap.

The Neandertal haplotype protective against severe COVID-19 is located in a region in chromosome 12 that contain the three genes *OAS1, OAS2* and *OAS3*, all of which encode oligoadenylate synthetases. These enzymes are induced by interferons and activated by double-stranded RNA. They produce short-chain polyadenylates, which in turn activate ribonuclease L, which degrades intracellular RNA, and activate other antiviral mechanisms in infected cells (reviewed by Choi *et al.* 2015).

To investigate which of these genes might be involved in the protection against severe COVID-19, we plot the genomic location of the *OAS* genes below the p-values for the association and the introgressed Neandertal haplotype (Fig. 1). The association (p<1e-6) overlaps all three *OAS* genes. The SNPs with the most significant associations (p<5e-8) are situated in the 3’-end of *OAS1* and in *OAS3*, and thus these two genes are most likely to be involved in the protective effect(s) of the Neandertal haplotype.

There are several SNPs on the Neandertal haplotype, which stand out as potentially functionally important in this region. One SNP (rs10774671) has been noted to affect a splice acceptor site (Sams *et al*. 2016). The most common allele at this SNP is derived and alters splicing of the *OAS1* transcript that produces an ancestral protein isoform (p46) to several alternative transcripts (Li *et al*. 2017). The ancestral isoform, preserved in Neanderthals, has been shown to have a higher enzymatic activity (Bonnevie-Nielsen *et al*. 2005). Outside Africa the ancestral allele is only present in individuals with the Neandertal haplotype whereas in Africa it exists independently of this haplotype presumably as a genetic variant inherited from the common ancestors of modern humans and Neandertals that was lost in the modern human population that left Africa (Sams *et al*. 2016). We note that a previous study (Carey *et al*. 2019) has identified recurrent *OAS1* loss-of-function mutations in primates, suggesting a cost of OAS1 enzymatic activity. In addition to the splice acceptor site, the 90kb Neandertal haplotype contains a missense variant (rs2660) in one *OAS1* transcript, a missense variant (rs1859330) and two synonymous variants (rs1859329, rs2285932) in *OAS3*, and a missense variant in *OAS2* (rs1293767).

The haplotype on chromosome 12 has previously been studied with respect to its effects on other viral infections. Notably, the Neandertal missense variant in *OAS1* (or variants in linkage disequilibrium with this variant) has been shown to be associated with moderate to strong protection against SARS-CoV (rs2660, OR = 0.42, 95% CI: 0.20-0.89) (He *et al*. 2006), although this study was limited in number of SARS-CoV cases and controls. The Neandertal-like splice acceptor variant has similarly been associated with protection against West Nile Virus (rs10774671, OR = 0.63, 95% CI: 0.5-0.83, Kim *et al*. 2009) and the Neandertal-like haplotypes have been associated with increased resistance to hepatitis C infections (El Awady *et al*. 2011). Furthermore, the Neandertal versions of the *OAS* genes are expressed differently in response to different viral infections both in terms of expression levels and splice forms (Sams *et al*. 2016). Thus, it is entirely possible that the *OAS* locus has been subject to distinct selective pressures by different pathogens during the hundreds of thousands of years that Neandertals and the ancestors of modern humans lived in western Eurasia and Sub-Saharan Africa, respectively.

After these groups came into contact and fused ~50,000 years ago, they became exposed to novel pathogens. Some of the variants contributed by Neandertals rose to high frequencies either locally, as is the case in South Asia for the Neandertal haplotype on chromosome 3 (Zeberg and Pääbo 2020), or almost everywhere in Eurasia, as is the case for the Neandertal haplotype on chromosome 12 (Fig. 2). With almost the entire world population now exposed to the novel SARS-CoV-2, the former haplotypes turn out to be detrimental while the latter haplotype turns out to be beneficial. Unfortunately, the protection against severe disease conferred by the Neandertal *OAS* locus is substantially smaller than the increased risk conferred by the chromosome 3 locus.

## Methods

The index variants for the three novel loci (rs10735079, rs2109069, rs2236757) were obtained from GenOMICC (Pairo-Castineira *et al.* 2020). The regional summary statistics from the round 4 (alpha) release of the meta-analysis carried out by the COVID-19 Host Genetics Initiative (https://covid19hg.org/results) was used to analyze the chromosome 12 locus. Linkage disequilibrium was calculated using LDlink 4.1 and alleles were compared to the archaic genomes using tabix (HTSlib 1.10). The Neandertal haplotype covering the region was investigated using sites where the Neandertal allele is not present in 108 Yoruba individuals (1000G Genomes Project), and the distances between the first two SNPs which cause the LD to fall below 0.75 in Europeans was taken as the length of the Neandertal haplotype. The probability of observing a haplotype of a certain length or longer due to incomplete lineage sorting was calculated as described (Huerta-Sánchez *et al.* 2014). The inferred ancestral states at variable positions among present-day humans were taken from Ensembl. Maps displaying allele frequencies of different populations were made using Mathematica 11.0 (Wolfram Research, Inc., Champaign, IL) and OpenStreetMap data.

## Acknowledgments

We are indebted to the COVID-19 Host Genetics Initiative (HGI) for making the GWAS data available and to the Max Planck Society and the NOMIS Foundation for funding.

## Supplementary material

**Table S1.** Genome-wide significant variants (p < 5e-8) on chromosome 12. Red marks the minor allele which in all cases shown is protective. Note that the human reference genome carries the Neandertal haplotype. *‘Ref’, ‘Alt’,* and ‘*Anc*’ refer to the reference, alternative and ancestral allele, respectively. LD denotes linkage disequilibrium in the 1000 Genomes Project dataset (r^2^).

| Chr | Pos | rsid | P-value | LD (rs4766664) | Ref | Alt | Anc | Vindija | Altai | Chagyrskaya | Denisova |
| --- | --- | --- | --- | --- | --- | --- | --- | --- | --- | --- | --- |
| 12 | 113362997 | rs4766664 | 5.7e-09 | 1.00 | T | G | T | T/T | T/T | T/T | G/G |
| 12 | 113381956 | rs2269899 | 7.9e-09 | 0.56 | C | T | T | C/C | C/C | C/C | T/T |
| 12 | 113380008 | rs10735079 | 7.9e-09 | 0.75 | G | A | A | G/G | G/G | G/G | A/A |
| 12 | 113410316 | rs10744791 | 1.3e-08 | 0.63 | G | A | A | G/G | A/G | G/G | A/A |
| 12 | 113372804 | rs4767037 | 1.4e-08 | 0.98 | A | C | A | A/A | A/A | A/A | C/C |
| 12 | 113372866 | rs1981557 | 1.4e-08 | 0.98 | G | C | G | G/G | G/G | G/G | C/C |
| 12 | 113372977 | rs1981555 | 1.4e-08 | 0.98 | G | A | G | . | . | . | . |
| 12 | 113376913 | rs7299132 | 1.5e-08 | 0.99 | T | A | A | T/T | T/T | T/T | A/A |
| 12 | 113357209 | rs1131476 | 1.6e-08 | 0.92 | G | A | G | G/G | G/G | G/G | A/A |
| 12 | 113357442 | rs2660 | 1.6e-08 | 0.92 | G | A | G | G/G | G/G | G/G | A/A |
| 12 | 113381695 | rs7131998 | 1.8e-08 | 0.86 | A | C | A | . | . | . | . |
| 12 | 113406196 | rs3937434 | 1.8e-08 | 0.63 | A | G | G | . | . | . | . |
| 12 | 113406460 | rs2016831 | 1.8e-08 | 0.63 | G | C | C | . | . | . | . |
| 12 | 113374017 | rs4767040 | 1.9e-08 | 0.98 | G | C | G | G/G | G/G | G/G | C/C |
| 12 | 113372961 | rs1981556 | 1.9e-08 | 1.00 | C | G | G | . | . | . | . |
| 12 | 113406945 | rs757405 | 1.9e-08 | 0.63 | T | A | A | T/T | A/T | T/T | A/A |
| 12 | 113373563 | rs3759376 | 1.9e-08 | 1.00 | A | G | G | A/A | A/A | A/A | G/G |
| 12 | 113409413 | rs4767045 | 1.9e-08 | 0.63 | C | T | T | T/T | T/T | C/C | T/T |
| 12 | 113402899 | rs4767044 | 2.0e-08 | 0.63 | C | A | A | C/C | C/C | C/C | A/A |
| 12 | 113373588 | rs3759375 | 2.0e-08 | 1.00 | A | G | G | A/A | A/A | A/A | G/G |
| 12 | 113386950 | rs2285932 | 2.0e-08 | 0.63 | T | C | C | T/T | T/T | T/T | C/C |
| 12 | 113375036 | rs7132797 | 2.1e-08 | 0.98 | A | C | A | A/A | A/A | A/A | C/C |
| 12 | 113392182 | rs7310667 | 2.1e-08 | 0.31 | A | G | A | . | . | . | . |
| 12 | 113381376 | rs6489882 | 2.2e-08 | 0.97 | G | A | A | A/A | A/A | A/A | A/A |
| 12 | 113377822 | rs6489879 | 2.3e-08 | 0.99 | G | A | G | G/G | G/G | G/G | A/A |
| 12 | 113379123 | rs7311182 | 2.3e-08 | 0.99 | T | C | C | . | . | . | . |
| 12 | 113405181 | rs1557866 | 2.4e-08 | 0.63 | A | C | C | A/A | A/A | A/A | C/C |
| 12 | 113380271 | rs6489880 | 2.4e-08 | 0.99 | C | T | T | . | . | . | . |
| 12 | 113364382 | rs4767033 | 2.4e-08 | 1.00 | T | A | A | T/T | T/T | T/T | A/A |
| 12 | 113378081 | rs4238033 | 2.4e-08 | 0.99 | T | A | A | T/T | T/T | T/T | A/A |
| 12 | 113375983 | rs1156361 | 2.6e-08 | 1.00 | T | C | C | T/T | T/T | T/T | C/C |
| 12 | 113372539 | rs7966314 | 3.1e-08 | 0.98 | A | G | A | A/A | A/A | A/A | G/G |
| 12 | 113408208 | rs2010604 | 3.2e-08 | 0.52 | G | C | C | G/G | G/G | G/G | G/G |
| 12 | 113374748 | rs10774679 | 3.3e-08 | 0.44 | C | T | C | C/C | C/C | C/C | C/C |
| 12 | 113385000 | rs10850103 | 3.3e-08 | 0.63 | A | T | T | A/A | T/A | A/A | T/T |
| 12 | 113376320 | rs3815178 | 3.3e-08 | 0.99 | C | T | C | C/C | C/C | C/C | T/T |
| 12 | 113376388 | rs1859330 | 3.4e-08 | 0.53 | G | A | G | G/G | G/G | G/G | A/A |
| 12 | 113381217 | rs6489881 | 3.6e-08 | 0.75 | A | T | A | A/A | A/A | A/A | T/T |
| 12 | 113380708 | rs7977345 | 3.7e-08 | 0.75 | A | T | T | A/A | A/A | A/A | T/T |
| 12 | 113379039 | rs7955267 | 3.7e-08 | 0.75 | C | T | C | . | . | . | . |
| 12 | 113382977 | rs2384074 | 3.7e-08 | 0.66 | C | T | C | . | . | . | . |
| 12 | 113396010 | rs10744789 | 4.1e-08 | 0.35 | T | C | C | . | . | . | . |
| 12 | 113371646 | rs9971885 | 4.2e-08 | 1.00 | A | C | C | . | . | . | . |
| 12 | 113378677 | rs4767041 | 4.3e-08 | 0.75 | G | A | G | . | . | . | . |
| 12 | 113367422 | rs2384072 | 4.6e-08 | 1.00 | T | C | T | T/T | T/T | T/T | C/C |
| 12 | 113366899 | rs7306205 | 4.6e-08 | 1.00 | A | G | A | A/A | A/A | A/A | G/G |
| 12 | 113367309 | rs1859336 | 4.7e-08 | 1.00 | C | T | C | C/C | C/C | C/C | T/T |
| 12 | 113364633 | rs4766667 | 4.8e-08 | 1.00 | G | A | G | . | . | . | . |
| 12 | 113370966 | rs1859333 | 4.9e-08 | 1.00 | T | C | C | T/T | T/T | T/T | C/C |
| 12 | 113368079 | rs6489877 | 4.9e-08 | 0.99 | A | G | A | . | . | . | . |
| 12 | 113363408 | rs6489866 | 4.9e-08 | 1.00 | A | G | A | . | . | . | . |

**Figure S1.**
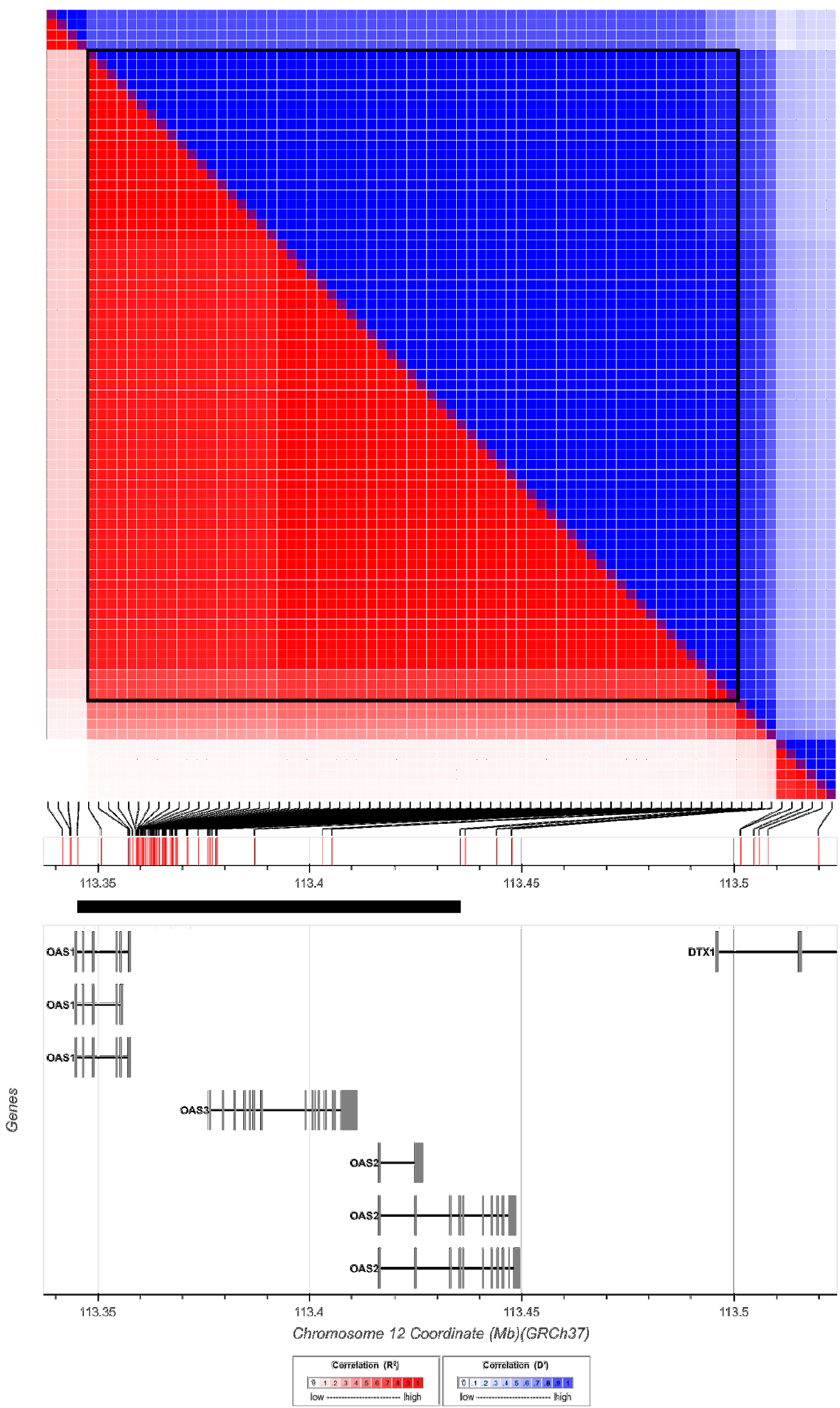
Diagnostic Neandertal variants and their pairwise linkage disequilibrium in Europeans. Black box indicates a set of 66 diagnostic variants in linkage disequilibrium (r^2^>0.75) defining a haplotype (black bar) with coordinates chr12:113,345,168-113,435,449 (*hg19*).

## Notes

### Competing Interest Statement

The authors have declared no competing interest.

### Summary of Updates

The revised manuscript use the index variants identified by the GenOMICC study (doi: 10.1101/2020.09.24.20200048), rather than using the meta-analysis of the COVID19 Host Genetics Initiative. No genome-wide data from the COVID19 Host Genetics Initiative are presented.

