## Supplementary figures and images for "A genetic variant protective against severe COVID-19 is inherited from Neandertals"

### Figure S1

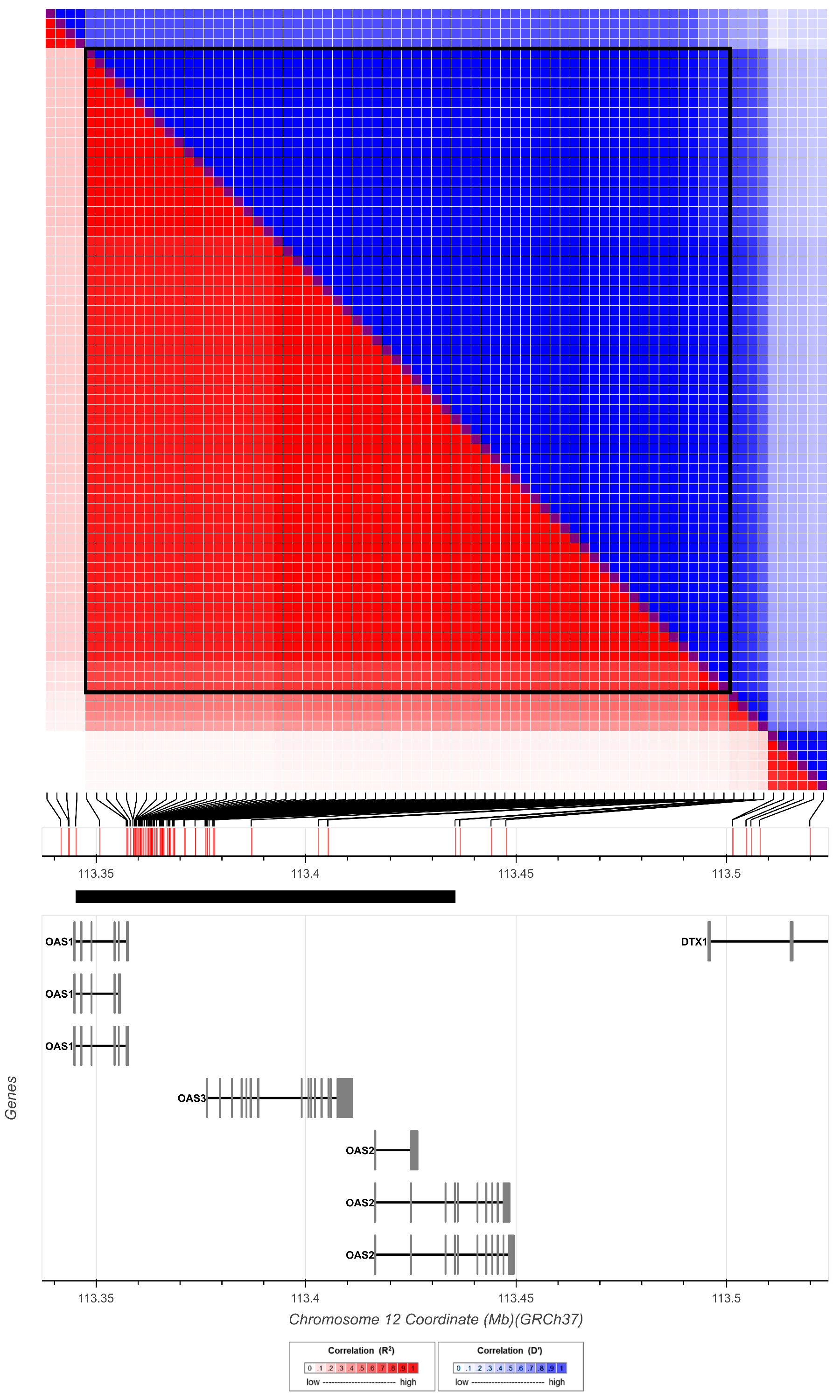

### Table S1

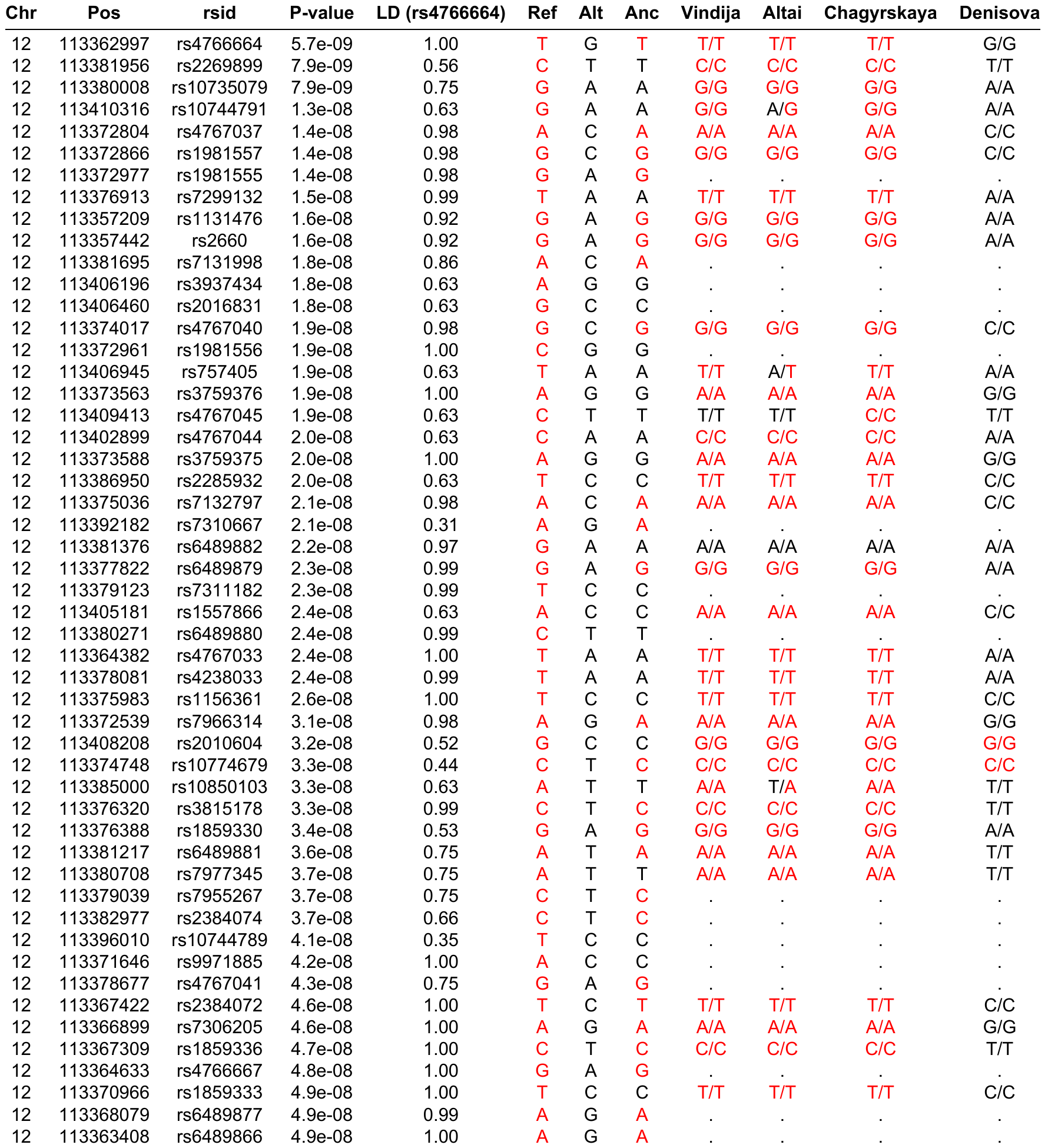
